# Spatiotemporal Gene Coexpression and Regulation in Mouse Cardiomyocytes of Early Cardiac Morphogenesis

**DOI:** 10.1101/349993

**Authors:** Yang Liu, Pengfei Lu, Yidong Wang, Bernice E. Morrow, Bin Zhou, Deyou Zheng

## Abstract

Cardiac looping is an early morphogenic process critical for the formation of four-chambered mammalian hearts. To study the roles of signaling pathways, transcription factors (TFs) and genetic networks in the process, we constructed gene co-expression networks and identified gene modules highly activated in individual cardiomyocytes (CMs) at multiple anatomical regions and developmental stages. Function analyses of the module genes uncovered major pathways important for spatiotemporal CM differentiation. Interestingly, about half of the pathways were highly active in cardiomyocytes at outflow tract (OFT) and atrioventricular canal (AVC), including many well-known signaling pathways for cardiac development and several newly identified ones. Most of the OFT-AVC pathways were predicted to be regulated by 6 6 transcription factors (TFs) actively expressed at the OFT-AVC locations, with the prediction supported by motif enrichment analysis of the TF targets, including 10 TFs that have not been previously associated with cardiac development, e.g., *Etv5*, *Rbpms,* and *Baz2b*. Finally, our study showed that the OFT-AVC TF targets were significantly enriched with genes associated with mouse heart developmental abnormalities and human congenital heart defects.

## Introduction

Prenatal heart development is controlled by evolutionarily conserved genetic networks (Olson, 2006, Waardenberg, Ramialison et al., 2014) consisting of well-recognized transcription factors (e.g., *Gata4*, *Nkx2-5* and *Tbx5*) (He, Gu et al., 2014, Takeuchi, Ohgi et al., 2003, Zhang, Nomura-Kitabayashi et al., 2014), signaling pathways (e.g., WNT and BMP) (Afouda, Martin et al., 2008, Wang, Green et al., 2014), and dynamic epigenetic networks that are modulated by histone modifications or DNA methylation (Chamberlain, Lin et al., 2014, Zhang, Wu et al., 2017, Zhang & Liu, 2015). These networks ensure the precise establishment of gene expression pattern and concurrent differentiation of cardiovascular cell types in an orderly spatial and temporal manner. However, transcriptional regulatory networks involved in cardiac looping have not been fully deciphered, partially due to the lack of appropriate experimental technology for resolving cellular and developmental heterogeneity.

The introduction of single cell RNA-seq technology has started to overcome this limitation. It revolutionizes the studies of gene regulation in embryonic developments by providing a systematic and high-throughput way to profile the expression of hundreds to thousands of cells simultaneously, resulting in the discovery of new cell types, cell state transitions and functions, as well as gene markers that are associated with unique cells populations and functions (Chen, Chakravarty et al., 2016, DeLaughter, Bick et al., 2016, Gladka, Molenaar et al., 2018, Li, Xu et al., 2016, Liu, Wang et al., 2017, See, Tan et al., 2017, Sereti, Nguyen et al., 2018, Skelly, Squiers et al., 2018). The scRNA-seq data, however, have generally not been fully exploited for addressing the dynamic expression of transcription factors and their cooperative or competitive interactions, even though it is well appreciated that the expression and targets of TFs are fundamental for deciphering the cardiac genetic programs.

In this study, we took a systematic approach to analyze single cell RNA-seq (scRNA-seq) data to characterize gene coexpression in differentiating cardiomyocytes at multiple anatomical locations of early cardiac developmental stages, and to investigate their regulation and maintenance by TFs. Our results indicate that the genetic programs for cardiac cell differentiation at the OFT-AVC are extremely complex, involving many critical pathways that are regulated by a significantly large number of TFs. This finding suggests that mutations in the genes regulating OFT-AVC development likely confers a high risk for congenital heart defects (CHD).

## Results

### Extraction of CM scRNA-seq data

The general approaches of our study are composed of construction of gene coexpression networks from scRNA-seq data, functional enrichment analysis, and prediction of genes and pathways regulated by transcription factors (Fig 1). Our study started from the scRNA-seq data that were previously obtained from mouse embryonic hearts at embryonic day (E) 8.5 to postnatal day (P) 21 (DeLaughter et al., 2016, Li et al., 2016). In these two studies, ~3,000 single cells in total were captured for RNA-seq from multiple dissected heart zones at different developmental stages. They were composed of multiple cardiac cell types, including endothelial cell (EC), mesenchymal cell (MC), and fibroblast-enriched cell (FE), and CM. To identify CMs from these data, we employed unsupervised clustering methods, similarly to the original reports. Principal component analysis (PCA) showed that cells from each of the two studies (“dataset 1” and “dataset 2”) could be segregated into two parts, confirmed by k-means clustering and t-Distributed Stochastic Neighbor Embedding (t-SNE) analysis (Fig. S1A-C). Based on the expression of known cardiac markers, we were able to identify the CM (expressing *Tnncl* and *Tnni3*), EC (expressing *Cdh5* and *Tek*) and MC (expressing *Collal* and *Ptn*) clusters (Fig. S1D). Although we were unable to directly compare our cell clustering result to the original reports because cell type information was not provided in the publically released data (DeLaughter et al., 2016, Li et al., 2016), the proportion of cells in these clusters and the expression profiles of known markers demonstrated that we have successfully reproduced the original clustering and successfully separated CM from other cardiac cells (Fig. S1D).

**Figure 1.**
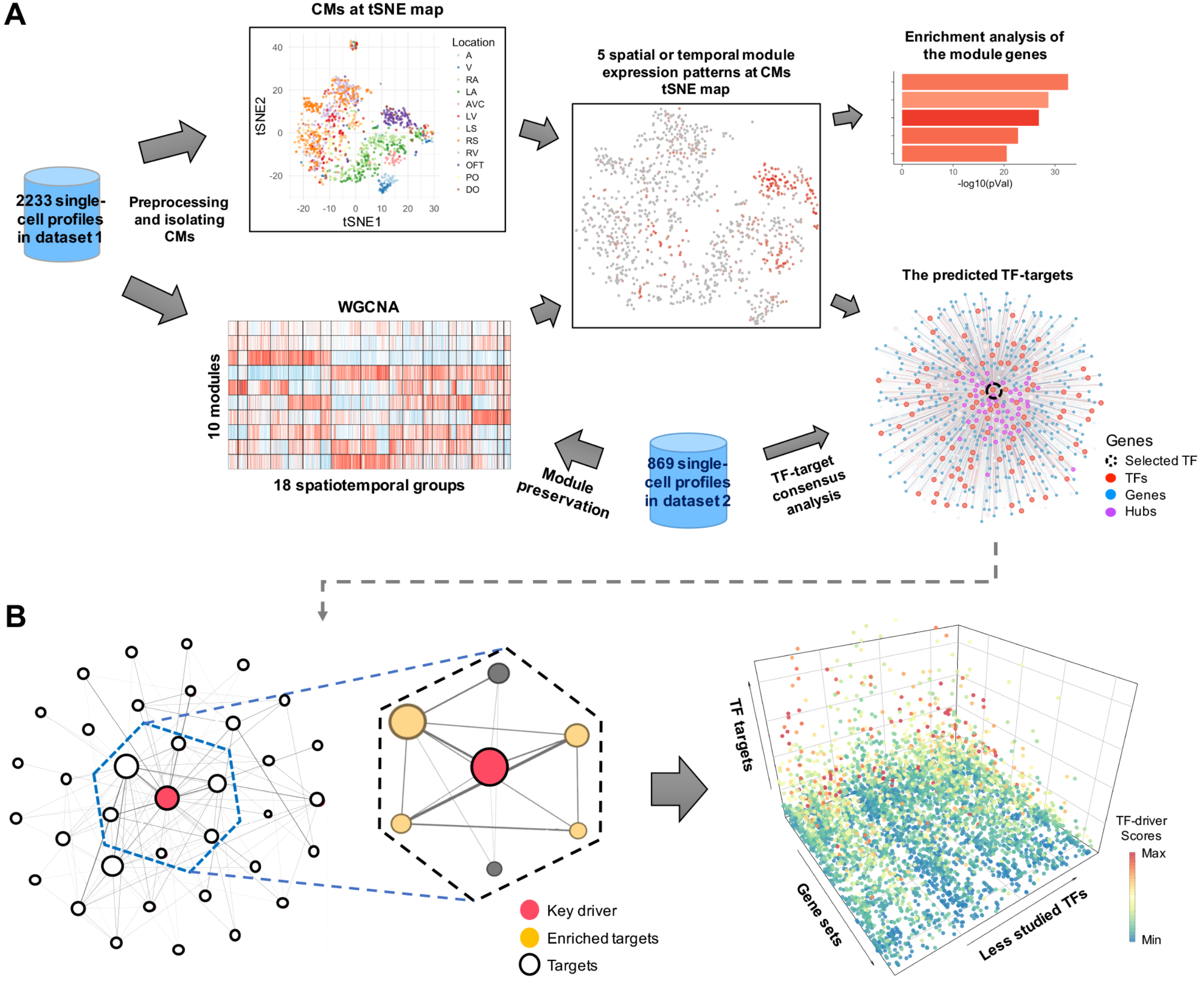
Workflow for inferring TF-target regulatory network. A Overview of our approach for identifying and characterizing gene co-expression patterns in CMs. The tSNE map indicates the gene expression similarity among CMs while WGCNA heatmap shows coexpression patterns of genes across CMs in different spatial or temporal subgroups. The plot for the predicted TF-targets displays a TF (dash circle) and its targets that are in the same gene module, with colors for different types of module genes. B Schematic illustration of function analysis for TF targets. Left panel shows that the potential functional roles of a TF could be predicted by enrichment analysis of its targets, with the dashed box highlighting targets of a TF (red). Right panel shows the enrichment results from the analysis of targets for less-studied TFs, with colors for TF-driver scores that quantify the potential regulatory effects of a TF on individual gene sets. See Methods for details.

### Spatiotemporal-specific transcriptional heterogeneity of CM

As CMs were captured at different developmental stages and anatomical locations, we first studied the contributions of these two factors (time and location) and other technical factors to gene expression variation (see Materials and Methods). We found that the number of expressed genes in individual CMs and anatomical origin explained the most expression variations, followed by developmental stage, but sequencing depth (“counts”) was not a main factor (Fig. S2A). Thus, we considered that the gene expression heterogeneity across CM groups mostly reflected biological difference, consistent with the original reports (DeLaughter et al., 2016, Li et al., 2016).

Accounting for developmental stages and anatomical locations, the CMs in dataset 1 could be approximately classified into four main clusters (Fig. 2A). Excluding outliers (cells in gray in Fig. 2A), E8.5 CMs appeared as one cluster, while the CMs at E9.5 and E10.5 could be segregated into three clusters based on anatomical locations: “atrial cluster” (including left atrium (LA), right atrium (RA) and AVC), “ventricular cluster” (including left ventricle (LV), left ventricular septum (LS), right ventricle (RV), right ventricular septum (RS)), and “outflow tract cluster” (including E9.5 outflow tract and distal outflow tract (DO)) (Fig. 2A and Fig. S2B). We also noticed that E10.5 CMs from proximal outflow tract (PO) showed higher gene expression similarity to RV CMs than E9.5 OFT or E10.5 DO CMs. In summary, the E8.5 to E10.5 CMs can be separated mainly by their temporal (developmental stages) and spatial (anatomic locations) origins that reflect their developmental trajectories.

**Figure 2.**
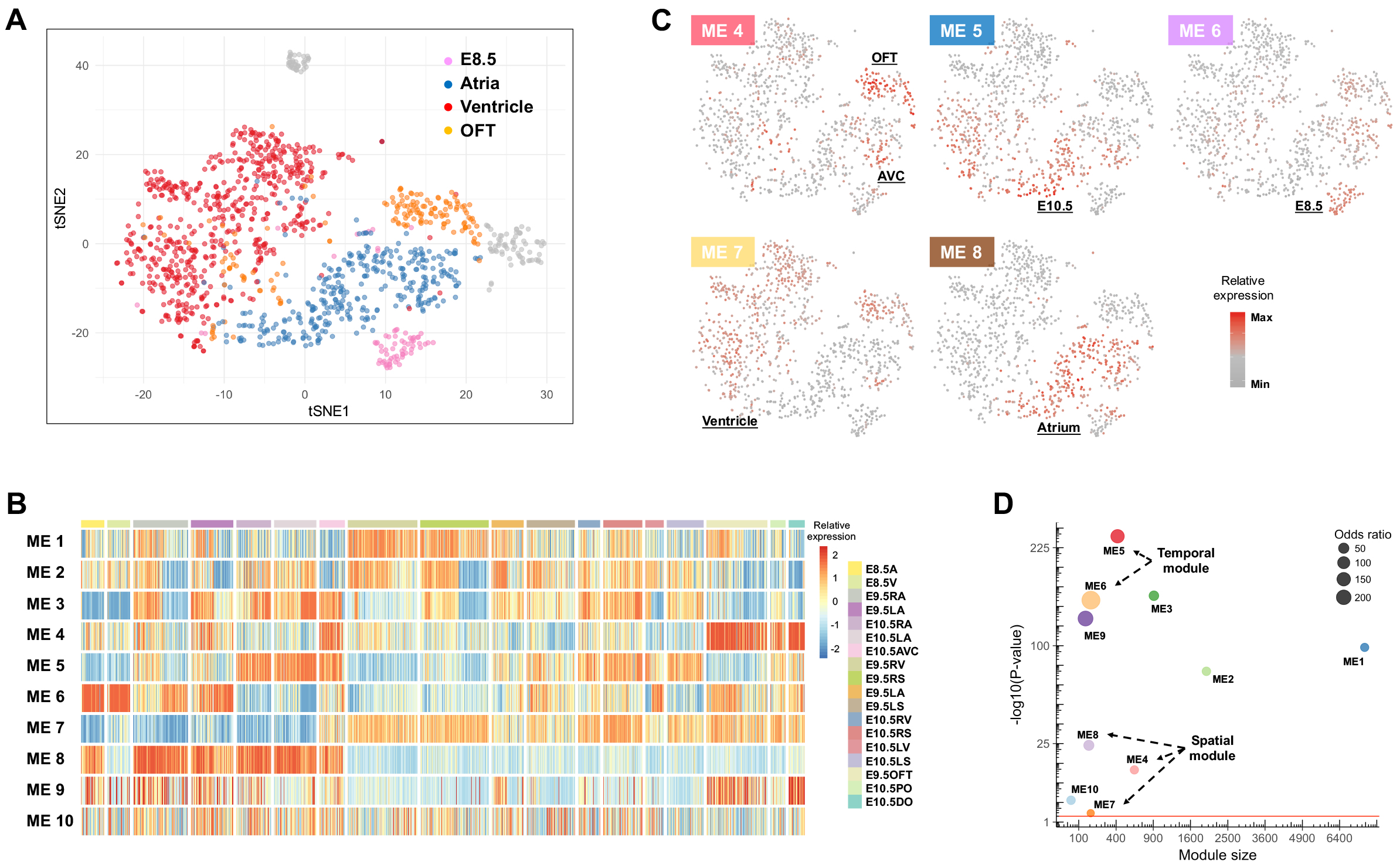
Identification of gene expression patterns for CMs at different anatomical-location or developmental-stage. A The t-SNE map of the CMs in dataset 1. B Ten gene modules generated by WGCNA. Columns of the heatmap represent individual CM cells, grouped by their anatomical and developmental stage origins, while each row is a gene module, with heat-map colors indicating the relative expression of its eigengene. C Module eigengene patterns at t-SNE map of CM cells, with colors for module eigengene expressions D Preservation analysis of the WGCNA modules. The dot size indicates the odds ratio (OR) and the y-axis shows the significance. The red line indicates the threshold (p-value < 0.05) for significant preservation.

We next carried out signed weighted gene co-expression network analysis (WGCNA, see Materials and Methods) to define co-expressed gene modules, and studied their relationship to the four main CM clusters. The analysis resulted in 10 gene modules from the 12, 792 genes expressed in both datasets (Fig 2B), with the module sizes ranging from 63 to 7,524. Interestingly, the 10 modules contained different proportions of noncoding genes and TFs (Fig S2C and Table S1), suggesting potentially different degree of regulation complexity. A comparison of the WGCNA result with the t-SNE cell clustering map and cell sample origins revealed that some of the module eigengenes, representing the first principal component of gene expression profiles in individual modules, displayed a strong correlation with specific CM clusters or groups (Fig 2B, C and Fig S2B). Therefore, our work unveiled CMs from different temporal or spatial origins for which we identified genes with highly similar expression patterns. To address the robustness of the gene modules, we performed consensus module analysis by incorporating the scRNA-seq data of CMs from E9.5 to 18.5 in dataset 2 (DeLaughter et al., 2016) (see Materials and Methods). The results confirmed that all gene modules were reproducible (i.e., “preserved”), indicating most modular genes have similar co-expression correlation in the two datasets, and the gene expression pattern is most likely a result of gene programs regulating embryonic CM development.

### Functional analysis of the WGCNA module genes

We performed GO (Gene Ontology) and pathway analyses to characterize the enriched functions of module genes (see Materials and Methods), focusing on the spatial and temporal modules (Fig 3A and Table S2). In the temporal modules, the genes of module 6 (E8.5 CM module) were enriched for function terms in regulation of cell differentiation, positive regulation of gene expression, and response to endogenous stimulus, including module genes *Gsk3b*, *Notch2* and *St13* (Fig 3B), indicating that this gene module is mainly involved in cell differentiation, concordant with the proliferation of early-stage CMs (de la Pompa & Epstein, 2012, Naito, Akazawa et al., 2005). The module 5 (E10.5 CM module) genes were enriched in oxidative phosphorylation, electron transport chain, and cardiac muscle contraction signaling known to be highly active in late-stage CMs (Chung, Dzeja et al., 2007). Not only did this module include tropoinin and ryanodine receptor genes, it but also contained a number of important electron transport genes from mitochondrial genome, such as *mt-Co2*, *mt-Co3* and *mt-Atp6* (Fig 3B), indicating distinct enhancement of mitochondrial ATP synthesis along with CM development (Table S2).

**Figure 3.**
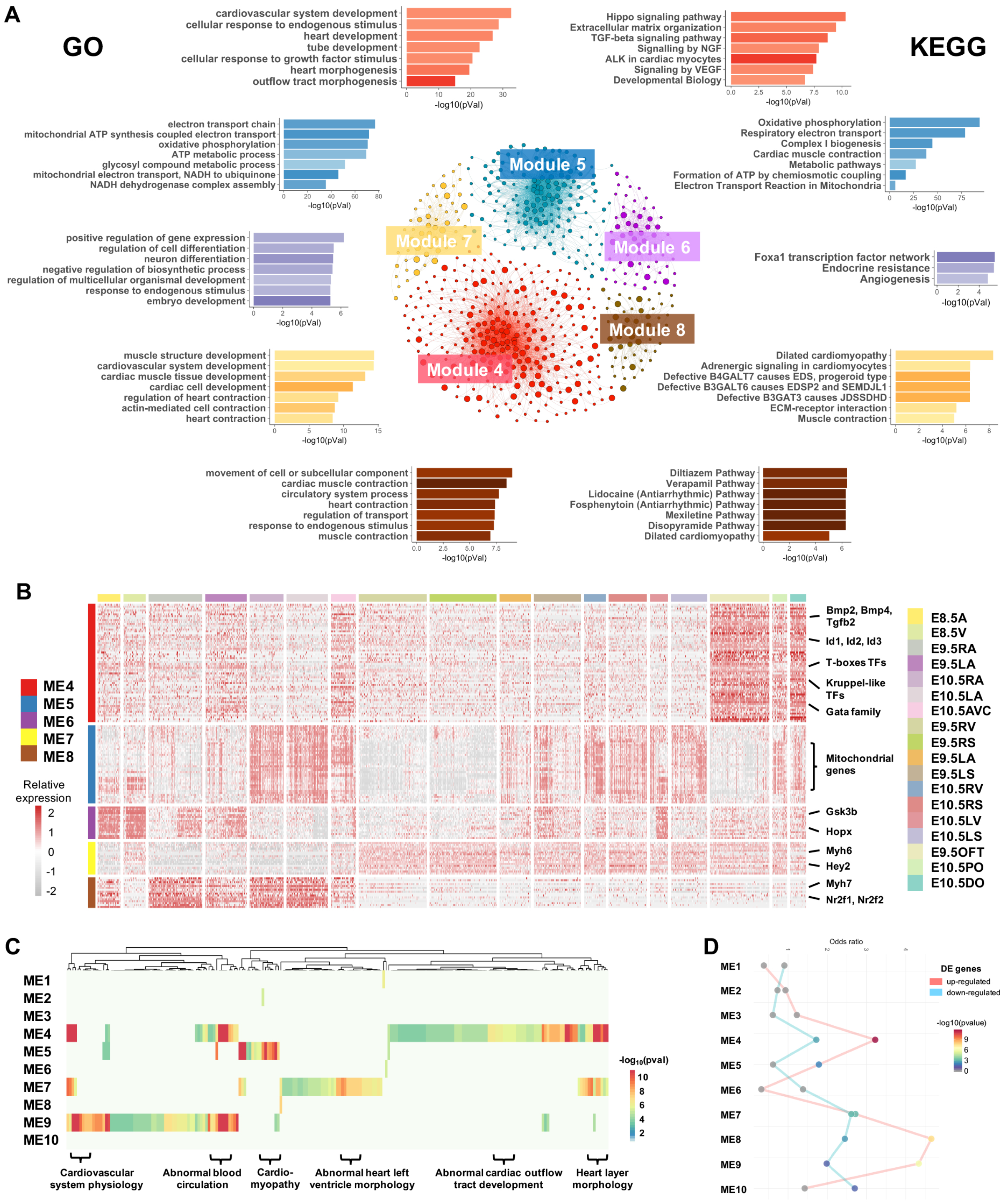
Enrichment analysis of genes in selected modules. A The top 7 representative enriched GO terms and KEGG pathways in the five selected modules. The networks in the middle illustrate the co-expression networks of genes in these modules, with the nodes representing genes and node sizes proportion to the numbers of co-expressed genes. Functional terms and pathways for individual modules are depicted by matching colors. B Expression patterns of hub genes in five selected modules. C Enriched phenotype- and disease-associated gene sets in the modules. The gene sets (columns) associated with heart phenotypes and diseases were enriched in the gene modules. The colors in the heat-map show statistical significance of enrichments, and some representative gene sets are labeled at the bottom. D Module enrichment of DE genes in dilated cardiomyopathy patients. The red and blue lines show the odds ratios of up-regulated and down-regulated genes, respectively. The dots with colors are statistically enriched (p-value < 0.05) while the gray ones are not.

In contrast, the OFT-AVC module genes (from spatial module 4) were enriched for terms important in cardiac morphogenesis, tube development, cardiovascular system development, and cellular response to growth factor stimulus. These processes have been reported to activate OFT remodeling (de la Pompa & Epstein, 2012). Interestingly, TGFβ, BMP and SMAD pathways, with their key regulators *Bmp2/4*, *Tgfb2*, *Id1/2/3* and *Smad6/7* (Fig 3B), were enriched in the module 4; they are known for roles in OFT and/or AVC development and morphogenesis (de la Pompa & Epstein, 2012, Hollnagel, Oehlmann et al., 1999, Wang et al., 2014). Interestingly, our analysis showed that genes of additional signaling pathways, such as NGF, VEGF, SCF-KIT, PDGF, MAPK, AKT, IGF, interleukins, leptin and insulin, were also significantly overrepresented in this module. These results indicate that OFT-AVC formation and remodeling are tightly regulated by a complex network of signaling pathways in response to both external growth factors and endogenous stimuluses. Our finding is consistent with previous studies of protein and gene networks critical for heart development (Lage, Greenway et al., 2012, Lage, Mollgard et al., 2010), but our analysis also uncovered additional novel pathways that were not described before, such as DAP12 (also called KARAP) signaling (Fig 3A and Table S2). For module 7 (ventricular CM module), function analysis identified enriched terms in muscle structure development, cardiovascular system development, and regulation of cardiac contraction. The module 8 genes, actively expressed in atrial CMs, exhibited significant enrichments in cell movement, response to endogenous stimulus, and ion signaling. Additionally, some well-known cardiac genes involving in cardiac contraction were members of these two modules, such as cardiac myosin *My12* and *Myh7* in module 7, and *Myh6* and tropomyosin *Tpm2* in module 8 (Fig 3B and Table S1). All these results underlined that our unbiased and systematic function analysis of spatiotemporal module genes uncovered both known and potentially novel signaling pathways in CMs at different anatomical locations or heart looping stages, thus providing molecular profiles for the diverse developmental processes and cellular functions in embryonic CMs.

### Modular enrichment of cardiac phenotype- and disease-associated genes

We next addressed how genes associated with different heart phenotypes or diseases were distributed in these gene modules, using the ToppGene and Fisher’s exact test (see Materials and Methods). Surprisingly, the results indicated that even though the modules 1, 2 and 3 contained >8 0% of the expressed genes, phenotype or disease marker genes were mostly enriched significantly in module 4, 5, 7 and 9 (adjusted p-value < 0.05) (Fig 3C and Table S3). In addition, reanalyzing the published transcriptional profiles of 218 dilated cardiomyopathy patients and controls (Liu, Morley et al., 2015), we found that 918 of the 1,661 differentially expressed genes (DE genes) were also expressed in the mouse scRNA-seq data, and they were statistically overrepresented in the modules 4, 5, 7, 8 and 9, concordant with our disease and pathway enrichment analyses above (Fig 3D and Table S2). These results support the value of our demarcation of gene modules, as it separated correctly the developmental and disease relevant gene modules from others. In addition, this analysis uncovered the genetic relationship between developmental process and heart abnormalities, thus highlighting the developmental bases of some heart diseases.

### Prediction of targets for spatiotemporal transcription factors

After studying the spatiotemporal expression of genes by co-expression networks and functional enrichment analysis, we set out to address the primary goal of current study, how the spatiotemporal expression and networks were regulated and maintained. In total, 186 TFs were present in the five selected modules (temporal- or spatial-specific modules). 103 of them were highly expressed in module 4 (OFT-AVC module), including T-boxes TFs (*Tbx2*, *Tbx3* and *Tbx20*), Kruppel-like TFs (*Klf3*, *Klf4*, *Klf6* and *Klf7*), Gata family (*Gata2*, *Gata3*, *Gata5* and *Gata6*) and Id family (*Id1*, *Id2* and *Id3*), as well as a large number of TFs whose functions in cardiac development remain unclear, like *Bach2*, *Etv5* and *Skil*. We also found that 12 TFs were in the E10.5 module (module 5), 2 7 TFs were active expression in E8.5 module (module 6), 20 and 24 TFs in ventricular module (module 7) or atrial module (module 8) respectively, including well studied cardiac genes *Hey2*, *Irx4*, *Sox5* and *Tbx5*. The expression patterns of some representative TFs in these modules were shown in Fig 4A.

**Figure 4.**
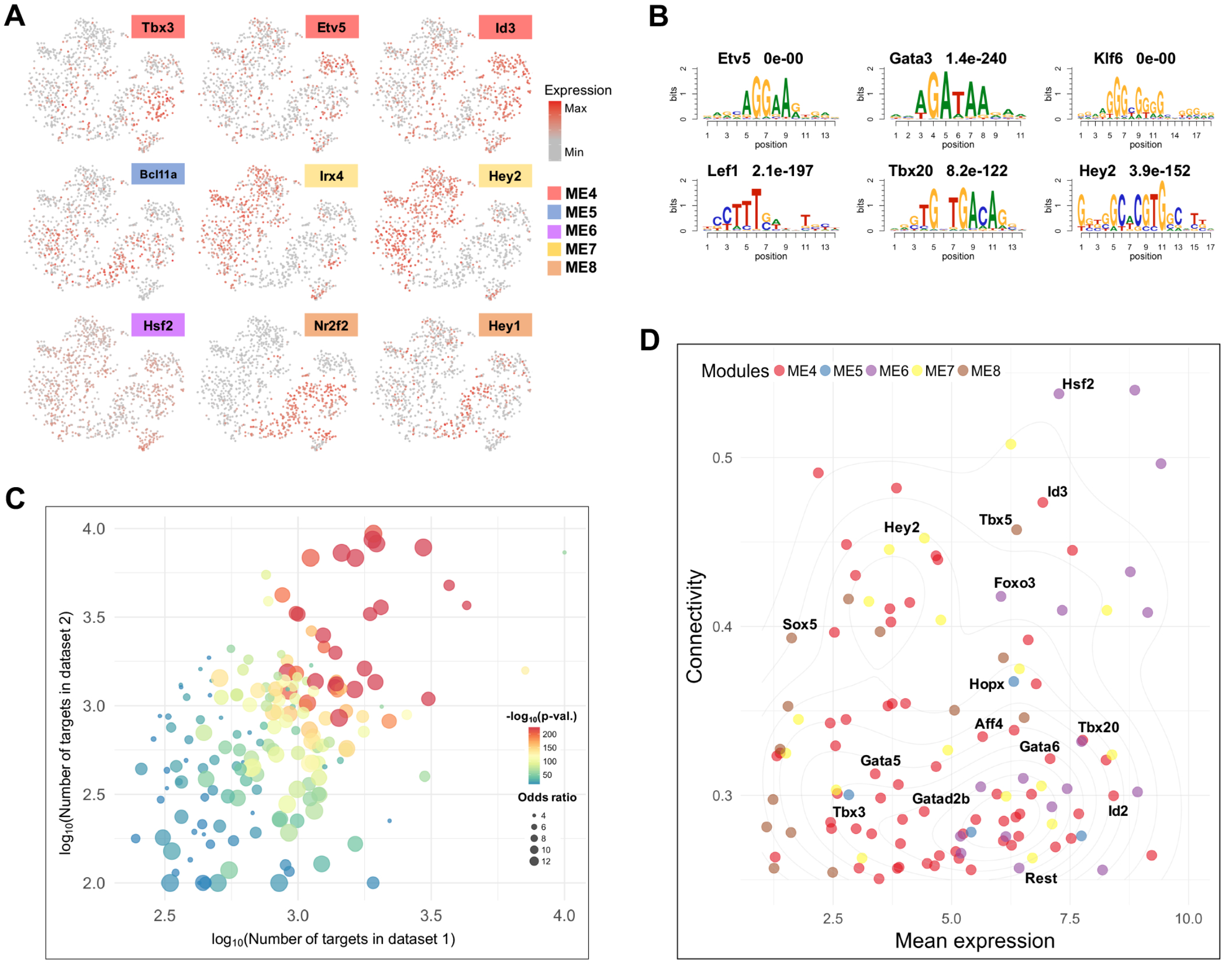
Spatiotemporally expressed TFs and their potential targeted genes. A Expression patterns of nine representative TFs in the selected modules. B The enriched motifs in the predicted targets of the representative TFs, with TF names and q-values at the top. C Preservation of TF-targets. Bubbles in the plot represent different TFs, with colors for the significance of preservation and sizes for odds ratios of target overlap. D Mean expression and eigengene-based connectivity of the candidate TF drivers (n = 123). In plot, x-axis represents the correlations between TFs’ expression profiles and their corresponding module eigengenes, while y-axis for average expression values for TFs.

To predict the targets for individual TFs, we used the adjacency scores computed by WGCNA for quantifying the co-expression strengths between genes, whereas considered a gene as a TF’s target if their pair-wise adjacency score was at the top 20% of all adjacency scores in the full gene co-expression network (see Materials and Methods). As such, TFs in the same module could have a very different list of targets. To support our TF-target prediction, we first performed motif analysis for 82 TFs that have known DNA binding motifs. The result showed that for ~80% of them, the corresponding motifs (or the motifs for the same TF family) were significantly enriched in the regulatory regions (defined by H3K27ac modification, see Materials and Methods) of their targets (Fig 4B and Table S4). Next, we re-analyzed several published microarray gene expression datasets, in which the expression of *Hey2*, *Nr2f2*, *Tbx3* and *Tbx5* in hearts was disrupted (Bonilla-Claudio, Wang et al., 2012, Luna-Zurita, Stirnimann et al., 2016, Russell, Ilg et al., 2015, Shirvani, Mookanamparambil et al., 2007, Wu, Kao et al., 2015). We asked whether the differentially expressed genes upon knockdown or knockout of these TFs were correlated to the targets predicted by our method, using the Chi-square test. The results showed significant correlations, Hey2 (p-value < 2.2e-16), *Nr2f2* (p-value = 1.246e-06), *Tbx3* (p-value = 0.00056) and *Tbx5* (p-value = 1.457e-05). Lastly, we repeated the TF-target prediction using the consensus gene network built by both scRNA-seq datasets and found that for 184 of the 186 TFs the predicted targets were reproducible (p-values < 0.05) (Fig 4C and Table S5, see Materials and Methods). To sum up, all these analyses and results support our TF-target prediction.

To study TFs with potentially major roles in cardiac-looping stage cardiomyocyte development, we then focused on TFs that are likely “key drivers”, central nodes in a co-expression network, (Carcamo-Orive, Hoffman et al., 2017, Langfelder & Horvath, 2008, Makinen, Civelek et al., 2014, Shu, Zhao et al., 2016, Zhang, Gaiteri et al., 2013) and studied what function pathways they may regulate. Specifically, we selected TFs that were hub genes and actively expressed in individual modules, resulting in 123 TFs (Fig 4D and Table S5, see Materials and Methods).

### Function analysis of well-known transcription factors across cardiac looping stages

From the 123 TFs, we started with 64 that were previously associated with cardiac phenotypes (referred to as “cardiac TFs”, see Materials and Methods), and asked in which pathways their targets were involved. Since we have already determined pathways, function terms and heart phenotype-associated gene sets enriched in each module, we only analyzed the enrichments of a TF’s targets in these pathways or terms. The analysis identified key drivers by “TF-driver scores,” which tested the association between a TF’s targets and an enriched pathway or term in a gene module (see Materials and Methods). As an example, shown in Fig 5A, TGFβ signaling pathway was enriched in the module 4 (Table S2), containing for 17 module 4 genes. Among them, 12 genes were predicted targets of *Id3* and 7 genes for *Pitx2*, while both *Id3* and *Pitx2* were in this signaling, with TF-driver score 1.28 and 2.84, respectively. This result suggested that these two TFs could be key drivers of this pathway (driver score > 1, see Materials and Methods), and that *Id3* was likely more important than *Pitx2* to the TGFβ signaling because of the former including more targets in this pathway, consistent with the previous report that *Id3* was directly activated by BMPs in TGFβ signaling (Hollnagel et al., 1999). Extending this TF driver analysis to all the 64 cardiac TFs, we found that ~80% (n = 51) of cardiac TFs were likely key regulators of the signaling pathways enriched in the five selected modules, with most cardiac TFs controlling multiple pathways (Fig 5B, Fig S3A and Table S6), such as the well-studied Gata family, T-boxes TFs, *G1i3*, *Hey1/2*, *Id1/3*, *Klf6*, *Loxl2*, *Mecom*, *Mef2c* and *Nr2f2*. On the other hand, ranking the pathways by the number of TF regulators, we found that regulations of the pathways in CM differentiation could be extremely complex, i.e. the top ranked pathways, like extracellular matrix organization, elastic fiber formation, TGFβ signaling, Hippo signaling, axon guidance and ALK in cardiac myocytes, were predicted to be regulated by >30 cardiac TFs simultaneously.

**Figure 5.**
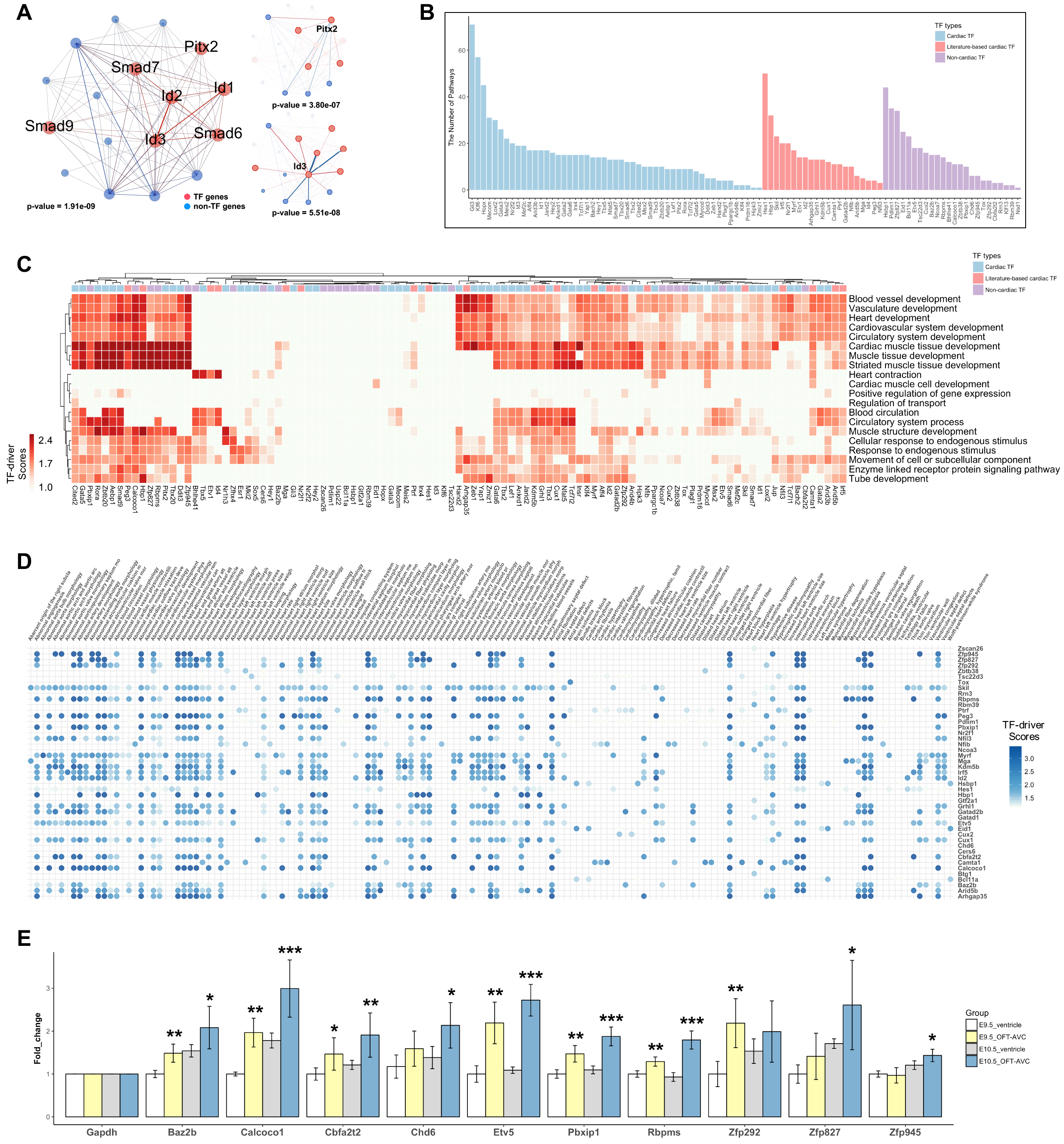
Function and phenotype analysis of cardiac and non-cardiac TF targets. A A network showing genes in the TGFβ signaling. In left panel, 17 genes of module 4 were in this pathway, with overrepresentation p-value = 1.91e-09. Node and edge connections in right panel represent adjacencies between a TF and its targets, 12 genes targeted by Id3 and 7 genes for Pitx2. B The number of enriched pathways (y-axis) under the regulations of cardiac, literature-based cardiac and non-cardiac TFs. C Predicted regulations of cardiac, literature-based cardiac and noncardiac TFs in 20 representative heart terms enriched in the selected WGCNA modules. The colors in heat-map shows the TF-driver scores in the functional terms, representing the regulatory effects of individual TFs on the terms. D Predicted functional roles of literature-based cardiac and noncardiac TFs in different heart phenotype- or disease-associated gene sets, with the colors for TF-driver scores. E qPCR analysis of 10 TFs highly expressed in OFT-AVC. The bar-plots show expression fold-changes (using Gapdh as reference). The statistical significances from comparisons of OFT-AVC to ventricle at E9.5 and E10.5 separately are indicated by “*”, “**” and “***” for p-values < 0.05, 0.005 and 0.0005, respectively.

We also extended this analysis to heart phenotype- or disease-associated gene sets. The results suggested that ~85% (n = 54) of cardiac TFs were involved in the regulation of gene sets implicated in multiple heart abnormality or diseases (Table S6), concordant with these TF being defined as “cardiac TFs”. Interestingly, the cardiomyopathy-associated genes were overrepresented in the modules 4, 5 and 7 (Table S2), and mainly regulated by *Gli3*, *Klf6*, *Loxl2*, *Hopx*, *Hey2*, *Irx4* and *Myocd*. In summary, our analysis, in an unsupervised manner, revealed the potentially functional roles of the spatiotemporally expressed cardiac TFs in differentiating CMs at critical cardiac-looping-stages.

### Analysis of targets for less studied transcription factors in heart looping

We next studied the 59 less-studied TFs (i.e., associations with abnormal heart phenotypes have not been clearly established), as analyses of their targeted pathways could provide new insights to their potential functions. Compared to the above results for cardiac TFs, the targets of these less-studied TFs, such as *Cux1*, *Etv5*, *Id2*, *Irf5*, *Klf4*, *Nfil3*, *Myrf* and *Skil*, were significantly enriched in many of the same heart developmental terms or processes, supported by similarly high TF-driver scores (Fig 5C and Fig S3B). Furthermore, our analysis showed that 45 of the 59 TFs were likely to regulate multiple functional pathways that were enriched in the selected gene modules (Fig 5B and Table S6), indicating their important functional roles. Interestingly, even though the mouse phenotype databases did not associate these TFs to an abnormal phenotype or disease, 21 of the 45 TFs have been reported to perform important functions in heart development, including *Hes1*, *Hbp1*, *Skil*, *Irf5*, *Id2*, *Cux1*, *Myrf*, *Kdm5b*, *Grhl1*, *Arhgap35*, *Gatad2b*, *Arid5b*, *Mga*, *Peg3*, *Nfil3*, *Nr2f1*, *Ptrf*, *Camta1*, *Etv1*, *Id4* and *Nfib* (Ahuja, Dogra et al., 2016, Albert, Schmitz et al., 2013, Chen, Qin et al., 2015, Cunnington, Nazari et al., 2009, da Rosa, Seminotti et al., 2016, Fletcher, Raza et al., 2015, Hirose-Yotsuya, Okamoto et al., 2015, Jongbloed, Vicente-Steijn et al., 2011, Loirand & Pacaud, 2014, Monnier, Iche-Torres et al., 2012, Muller-Borer, Esch et al., 2012, Nassiri, Liu et al., 2010, Qi, Yu et al., 2017, Quaranta, Fell et al., 2018, Rochais, Dandonneau et al., 2009, Seneviratne, Edsfeldt et al., 2017, Shekhar, Lin et al., 2016, Song, Gao et al., 2018, Taniguchi, Maruyama et al., 2016, Theis, Sharpe et al., 2011, Waldron, Steimle et al., 2016, Walkowska, Pawlak et al., 2016, Yaniz-Galende, Roux et al., 2017). As such, we named these TFs as “literature-based cardiac TFs”, to distinguish them from the other “non-cardiac TFs” that do not have known evidence for a cardiac function (Table S5). These findings support the biological relevance of our TF-target prediction, and strongly indicates that the set of non-cardiac TFs (n = 24) still have high possibilities to regulate critical pathways in embryonic heart development.

Performing enrichment analysis of TFs’ targets for the heart phenotype- or disease-associated gene sets enriched in the five spatiotemporal modules, we found that the targets of the 21 literature-based cardiac or the 24 non-cardiac TFs were significantly overrepresented in multiple gene sets, such as abnormal cardiac outflow tract development, abnormal cardiovascular development, abnormal heart ventricle morphology, and abnormal cardiac muscle contractility (Fig 5D and Fig S3C-D). Notably, >60 phenotype- and disease-associated gene sets were found to be under the regulations of *Hes1*, *Skil*, *Myrf*, *Etv5*, *Cux1*, *Irf5* and *Id2*. Most of them, except *Etv5*, have been reported to play a functional role in heart development, suggesting the potential heart genotype-phenotype associations of these TFs.

To confirm the high expression of the OFT-AVC TFs, we selected 10 less studied TFs for qPCR analysis. The results showed that almost all of these 10 TFs had significantly higher expressions at OFT-AVC than ventricles at E10.5, with some also exhibiting higher expression in E9.5 OFT (Fig 5E). Further analysis found that the more specific expression patterns they had, the more heart-phenotype gene sets these TFs are involved in, with spearman correlation > 0.7 and p-value < 0.05 at both E9.5 and 10.5 stages (Table S6).

### Expression changes and functional roles of transcription factors in subgroups of atrial or ventricular CMs

The above analyses considered CMs from the same spatial or temporal origin as a “homogenous” group, but CMs in the same group actually showed subtle transcriptional heterogeneity, because CMs from the same anatomical location could be a mixture of cells at multiple developmental states. Therefore, we constructed putative developmental trajectories based on top variable genes (see Materials and Methods) in order to study potential cell state transitions of CM development at atrium, ventricle or outflow tract (including E9.5 OFT, E10.5 DO and PO) (Fig 6A-B and Fig S4A). The analysis revealed three distinct subgroups (“A”, “B” and “C”) in each of the trajectories, largely reflecting the temporal and spatial relationship of CMs at those locations (Fig S4B-G).

**Figure 6.**
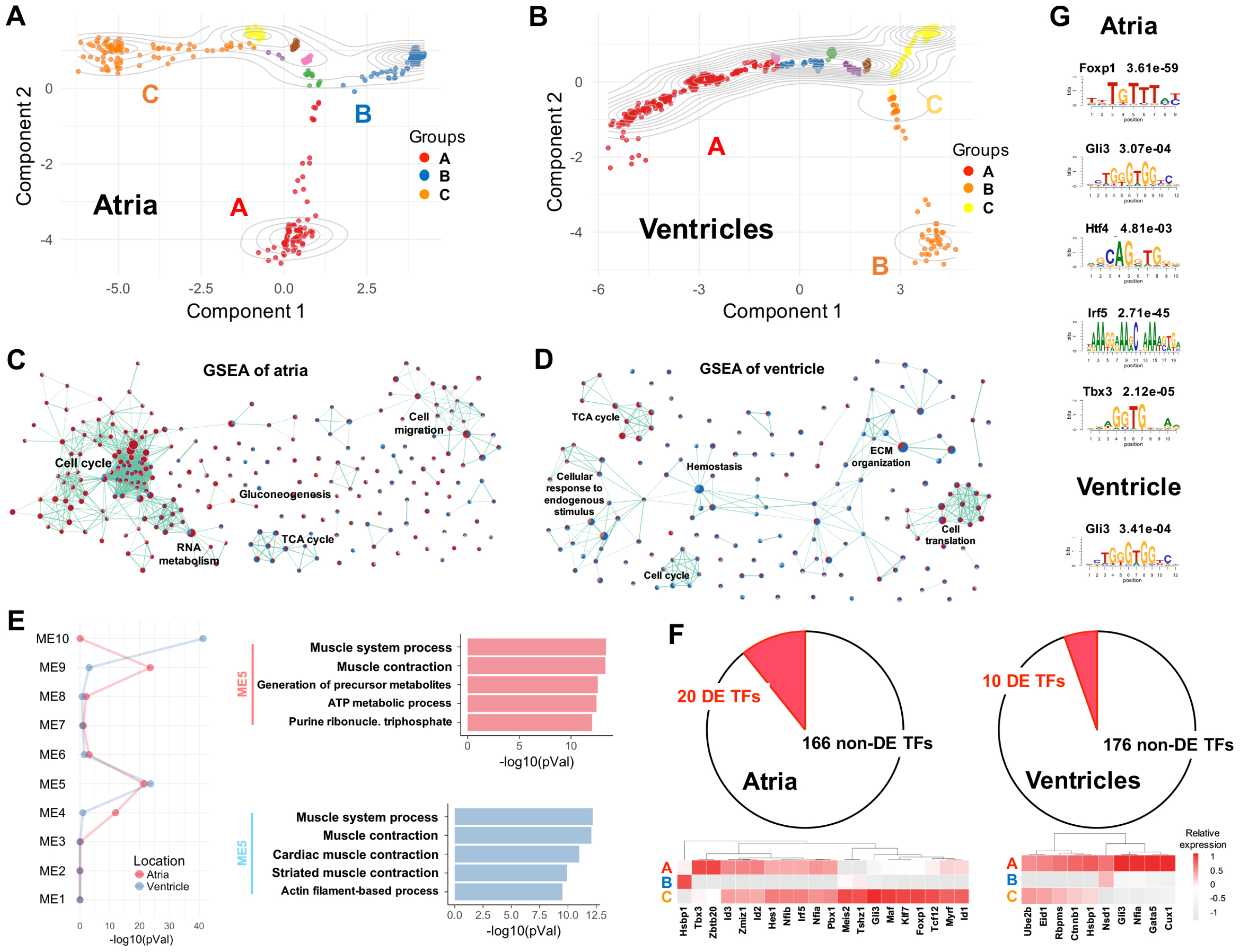
Differentially expressed genes and TFs in atrial or ventricular CMs. A A trajectory of atrial CMs, indicating that the atrial CMs are consisted of three major subgroups (A, B and C). B The trajectory of ventricular CMs. C - D Enriched gene sets for the DE genes among the three subgroups of atrial (C) and ventricular (D) CMs. Here, DE genes from each of the pair-wise comparisons were used for GSEA. The up-regulated and down-regulated genes are in red and blue nodes, respectively, while the edges of nodes represent the overlapping genes in two pathways. E Module enrichments of atrial and ventricular DE genes. The enrichment significance of atrial or ventricular DE genes were represented by the red and blue lines, respectively in left panel. The right panel shows the top 5 enriched pathways of atrial or ventricular DE genes that were also identified in module 5. F Spatiotemporally DE TFs at atria and ventricles. G The enriched motifs for atrial or ventricular DE TFs in panel (F).

Gene set enrichment analysis (GSEA) indicated that DE genes from pairwise comparisons of the three atrial subgroups were enriched for developmental pathways, such as cell cycle, mRNA processing, protein metabolism, ECM organization, citric acid cycle and gluconeogenesis (Fig 6C and Table S7). In comparison, the enrichment of ventricular DE genes among the three subgroups was mainly related to myogenesis, oxidative phosphorylation, angiogenesis, epithelial mesenchymal transition, apical junction, and Heme metabolism pathways (Fig 6D and Table S7). Interestingly, only a small number of DE genes (< 60) were found among the subgroups of OFT CMs, even though CMs at this anatomical location were from two embryonic heart stages (E9.5 and E10.5). Mapping the DE genes to the above module genes showed that atrial DE genes were significantly enriched in the modules 4, 5 and 9, while ventricular DE genes were enriched in the module 5 and 10 (Fig 6E). Pathway analysis of the overlapped DE genes suggested that both atrial and ventricular DE genes were dramatically overrepresented in the module 5 and enriched in pathways related to muscle system process and muscle contraction (Fig 6E), indicating our trajectories could represent developmental states of CMs at different anatomical heart regions.

To further support the importance of TFs in CM development, we examined TFs that were differentially expressed in atrial or ventricular subgroups, and included in the five gene modules. This resulted in 20 DE TFs in the atrial and 10 in the ventricular subgroups (Fig 6F and Fig S4D). Motif analyses showed that the regulatory regions of these atrial DE genes were enriched for 5 TF motifs, *Foxp1* (q-value 3.61e-59), *Gli3* (q-value 3.07e-04), *Htf4* (q-value 4.81e-03), *Irf5* (q-value 2.71e-45) and *Tbx3* (q-value 2.12e-05) (Fig 6G). For the ventricular DE genes, *Gli3* motif was found (q-value 3.41e-04) (Fig 6G). The findings strongly supported the regulations of CM differentiation by these TFs that are expressed in a highly spatiotemporal manner.

## Discussion

Our study defined spatiotemporal gene co-expression networks from cardiac scRNA-seq data, examined their enrichment of functional pathways and developmental phenotypes, but more importantly investigated TFs’ roles in regulating or maintaining functional gene modules that are active in CMs at a specific anatomical location or developmental stage during heart looping stages, from E9.5 to 10.5. Our findings not only confirmed the critical functions of 51 well-known cardiac TFs, like T-boxes gene family, Kruppel-like gene family, Id gene family, and Gata gene family, but also uncovered the potential novel functional roles of 45 less studied TFs. The importance of the latter group of TFs in regulating CM developmental signaling and pathways is supported by the significant enrichment of their targets in heart development and phenotypes (Fig 7).

**Figure 7.**
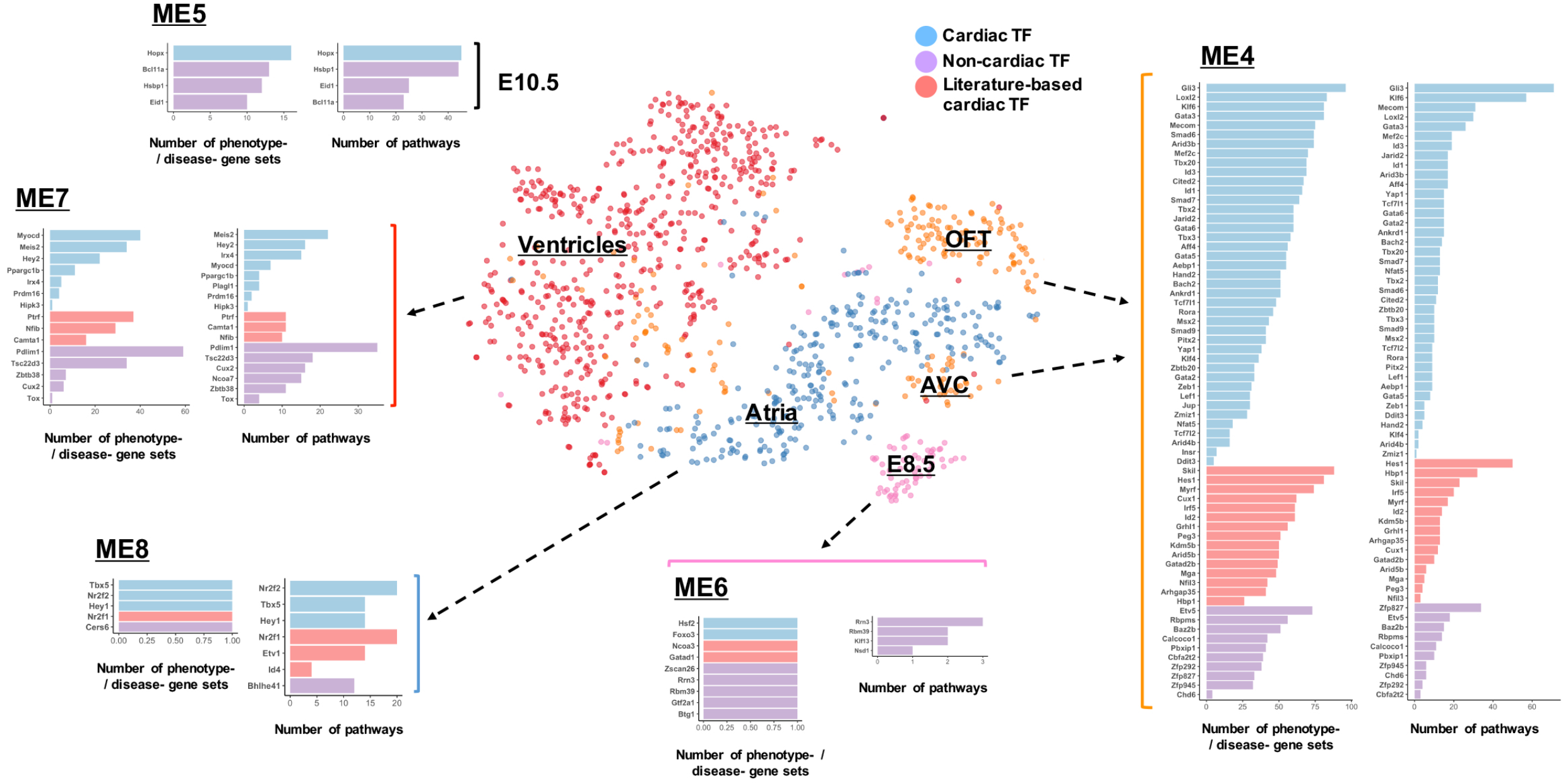
Summary of TFs and their regulatory functions in the spatiotemporal modules. The functional roles of the candidate TFs, including cardiac, literature-based cardiac and non-cardiac, in the five selected modules. Each bar-plot panel of individual modules shows the number of the enriched functional pathways and heart phenotype- or disease-associated gene sets under the regulations of the designated TFs.

Our analysis showed that across heart looping stages many signaling and pathways involved in CM differentiation were enriched in the OFT/AVC module (module 4) and they were predicted to be highly regulated by multiple TFs, and more so than genes in the other modules (Fig 7). In addition, we found that the proportion of TFs in this module was much higher than those of other modules (Fig S2C). Integrating the results of function analysis of “cardiac”, “literature-based cardiac” and “noncardiac” TFs, our study strongly suggests that the genetic program in differentiating CMs at OFT/AVC is not only regulated by well-studied cardiac TFs, like *Tbx2/3*, but also under the complex regulation of other 64 TFs (Fig 7), such as *Gli3*, *Mecom*, *Loxl2* and *Meis2*, including additional TFs whose cardiac functions are yet to be addressed. The high expression of the 10 non-cardiac TFs in OFT/AVC was confirmed by qPCR analysis (Fig 5E). Interestingly, we also found a significant correlation between the non-cardiac TFs’ expression patterns and the number of heart phenotypes they involved, with greater specific spatial expression correlated with more phenotypes. Considering all the results described here, we hypothesize that OFT/AVC formation and remodeling is likely the most tightly regulated embryonic heart developmental process. As such, mutations in the genes, especially TFs, important for OFT/AVC development are likely to be the high-risk factors for congenital heart defects. In fact, we analyzed two sets of manually curated CHD-associated genes and found that they were significantly enriched in the OFT-AVC module, with the odds ratios of 2.39 (p-value = 0.001) and 2.35 (p-value = 0.0003) for the 186 CHD genes in the Sifrim et al (Sifrim, Hitz et al., 2016) and the 253 CHD genes in the Jin et al (Jin, Homsy et al., 2017), respectively. We also analyzed the genes with significantly higher expression in ECs and MCs of OFT-AVC than the ECs/MCs of other locations. The results indicated that only the group of genes with higher expression in OFT-AVC ECs were significantly enriched for the 253 CHD genes curated by Jin et al, suggesting cell type specificity of our finding.

Although our study has analyzed more than one thousand cardiac CMs from multiple developmental stages and anatomical locations, the number of total cells is relatively small, considering that new scRNA-seq technology allows expression profiling of tens of thousands or even up to millions of cells (Zheng, Terry et al., 2017). The inclusion of more cells from more time points can certainly improve our results, especially in terms of finding key TFs regulating cell fate commitment and lineage specification. In theory, with all cardiac cells from the same heart being sampled simultaneously, one can also study the TF-regulatory networks and signaling cross-talks among different cardiac cell types, an important matter not addressed in the current study. Finally, it will be necessary to validate our TF-target predictions using scRNA-seq data from wild type hearts and mutant hearts with the expression of selected TFs compromised, such as previous studies of Nkx2.5 function (DeLaughter et al., 2016, Li et al., 2016). Nevertheless, the current study has identified key TFs, including not fully characterized ones, that can play important regulatory roles in embryonic heart development.

## Materials and Methods

### Single cell RNA-seq data

Single cell transcriptional profiles of murine embryonic cardiac cells (dataset 1) was obtained from the GEO (GSE76118). The cells were captured from different anatomical locations at different time points and contained a total number of 2,233 cells: CMs (1252), EPs (40), ECs (191), MCs (281), and the cells with two or more cell type characters (360) (Li et al., 2016). Dataset 2 (DeLaughter et al., 2016) was downloaded from the GNomEx database, with accession numbers 272R, 274R, 275R, 276R, 277R1, 279R1, 280R, 281R, 439R (https://b2b.hci.utah.edu/gnomex/). The 869 cells were mainly isolated from atrium or ventricle of murine heart cells from embryonic to postnatal days, including 484 CMs, 146 EDs, 104 FBs and 135 other cells.

### Data processing and quality control

Raw counts of gene expression data in dataset 1 were directly obtained from the GEO, with read aligned by STAR (2.4) and expression qualified by HTSeq (0.6.1), according to the original authors (Li et al., 2016). For dataset 2, only fastq files were available in the GNomEx database. Thus, HISAT2 (Kim, Langmead et al., 2015) was employed to align reads to the mouse genome (GRCm3 8.79), followed by SAMtool 1.4 (Li, Handsaker et al., 2009) to generate alignment files and HTSeq (0.6.1) (Anders, Pyl et al., 2015) to compute read counts. In dataset 1, read counts per cell ranged from 0.55 to 3.3 million, and the number of expressed genes were >4,000. Cells not meeting these criteria were eliminated for subsequent study. Similarly, cells in the dataset 2 were retained if the counts were 0.3 to 2.1 million and >2500 genes were expressed. Furthermore, genes with reads in less than 50% of cells were discarded, resulting 12,792 genes for further analysis, including 1,275 TF genes, 9,694 protein-coding genes, and 1,823 non-coding genes. In both datasets, cells with two or more cell types were deleted, based on our clustering analysis.

### Cell clustering, dimension reduction, and trajectory construction

The raw gene count data were processed using packages tidyr 0.8.0 (Wickham & Henry, 2018), dplyr 0.7.4 (Wickham, Francois et al., 2017) and scater 1.6.1 (McCarthy, Campbell et al., 2017) in R. Clustering cells was carried out mainly with principle component analysis (PCA), t-distributed stochastic neighbor embedding (t-SNE) by package Rtsne 0.13 (van der Maaten & Hinton, 2008), and k-means clustering by package SC3 1.6 (Kiselev, Kirschner et al., 2017) in R. Ordering cells in a trajectory corresponding to a biological process was also performed by the package monocle 2.6.1 (Trapnell, Cacchiarelli et al., 2014). We carried this analysis for atrial CMs, using data from LA, RA and AVC of E9.5 and 10.5, ventricular CMs, using data from LS, RS, LV and RV of E9.5 and 10.5, and OFT CMs, using data from OFT, PO and DO. In each case, the top 1,000 most dispersed genes were used in an unsupervised manner. Further, plots in this study were generated by ggplot2 2.2.1 (Wickham, 2016) while heat-map built by pheatmap 1.0.8 (Kolde, 2013) in R.

### WGCNA co-expression network

Signed weighted gene co-expression network was built from the expressed genes in dataset 1, using the package WGCNA 1.61 (Langfelder & Horvath, 2008) in R, with the scale-free topology fit index over 0.8. More specifically, we estimated soft-thresholding power and chose 10 to construct gene network in dataset 1. In WGCNA, the adjacencies are used to represent gene connection strengths of gene network. In signed weighted network,

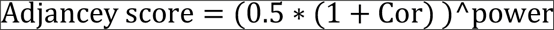

Cor: correlation coefficient, power: soft-thresholding power

From the intramodular connectivity analysis, we defined hub genes as the top connected genes (eigengene-based connectivity, or called signed connectivity, between the gene and module eigengene (kME) ≥ 0.25).

### Consensus module analysis

The preservation of the gene modules in dataset 1 were supported by consensus module analysis using the package WGCNA 1.61 (Langfelder, Luo et al., 2011). The analysis built consensus modules by combining dataset 1 and dataset 2 and then test if the modules generated with dataset 1 alone were preserved in the consensus modules, i.e., if the genes of a given module in dataset 1 were significantly enriched in any consensus module from both datasets, with p-value < 0.05. The Cytoscape (Shannon, Markiel et al., 2003) and Gephi tools (Bastian, Heymann et al., 2009) were employed to visualize gene co-expression networks of the modules; the node size represented connections and the length of edge indicated the strength of the connection.

### Differential expression analyses

The differentially expressed genes of different single-cell clusters or subgroups were identified by using package edgeR 3.20.1 (Robinson, McCarthy et al., 2010) in R, with the adjusted p-value < 0.05 as significant. Batch effects were removed when detecting differentially expressed genes, including spatial factor, temporal factor and the condition variable.

### Microarray data and DE analysis

The microarray of cardiomyopathy patients were obtained from the GEO (GSE57338) (Liu et al., 2015), and the data for Bmp2/4, Hey2, Nr2f2, Tbx3 and Tbx5 were also downloaded from the GEO, with accession numbers GSE34502, GSE6526, GSE63759, GSE73862 and GSE77576, respectively (Bonilla-Claudio et al., 2012, Luna-Zurita et al., 2016, Russell et al., 2015, Shirvani et al., 2007, Wu et al., 2015). All the microarray data were processed using the package oligo 1.42.0 (Carvalho & Irizarry, 2010) in R, and the DE genes was detected using package limma 3.34.5 (Ritchie, Phipson et al., 2015) in R, with adjusted p-value < 0.05.

### Gene sets, phenotype and disease-associated enrichment analyses

Pathway (GO, KEGG and Reactome) (Fabregat, Jupe et al., 2018, Kanehisa, Furumichi et al., 2017, The Gene Ontology, 2017) and gene set enrichment analyses were performed using the ToppGene suite (Chen, Bardes et al., 2008) and GSEA 3.0 (Subramanian, Tamayo et al., 2005), respectively, with FDR < 0.05 as a cutoff in David and FDR < 0.1 in GSEA. Phenotype and disease-associated enrichment analyses were also carried out using the ToppGene suite, with FDR < 0.05. The enriched phenotype and disease terms (IDs) from the ToppGene were cross compared to the disease category information in the MGI (Mouse Genome Informatics) (Dickinson, Flenniken et al., 2016), IMPC (International Mouse Phenotyping Consortium) (Collins, Finnell et al., 2007) and DisGeNET (Pinero, Bravo et al., 2017) databases to retrieve terms related to cardiovascular systems, heart phenotypes and cardiac diseases, resulting in 395 gene sets associated with abnormal heart phenotypes and 99 gene sets linked to cardiac diseases.

The MGI: http://www.mousemine.org/mousemine/begin.do
The IMPC: http://www.mousephenotype.org/data/search/mp?kw=*
The DisGeNET: http://www.disgenet.org/web/DisGeNET/menu/browser?1.

### Cardiac, literature-based and non-cardiac TFs

To uncover potentially novel TFs involved in regulating cardiac development, we used heart phenotype- or disease-associated gene sets from the MGI, IMPC and DisGeNET databases, and literature search to separate TFs into three groups. “Cardiac TFs” were the TFs that have been associated (i.e., annotated) within any heart-phenotype gene sets from the three databases. The rest were further classified either as “literature-based cardiac TFs,” if there is a literature support for a function in heart development, or otherwise as “non-cardiac TFs”, to indicate their unknown roles in cardiac development or function.

### TF-target analysis

123 TFs considered as key drivers were selected by these criteria, 1) a member of the five selected modules (4, 5, 6, 7, 8), 2) high expression (average cpm > 1 in the given module), and 3) a hub gene in the module. To predict the TF-target relationship, we used the adjacency scores computed by WGCNA. Considered all the genes (n = 12,792) in the signed weighted gene co-expression network of dataset 1, the top 20% adjacency values were ≥ 0.0017. For a TF, any gene with an adjacency ≥ 0.0017 was considered to be a candidate target of this TF. In TF-target preservation analysis, this cutoff (adjacency ≥ 0.0017) was also applied to define TF-targets in consensus gene co-expression networks from both datasets. Because TF-target analysis was based on the top adjacency, a TF and its targets might be in different modules, and our TF-target consensus analysis included all the targets. On the other hand, to investigate the regulatory roles of TFs in the spatiotemporal modules, our enrichment analysis of a TF’s targets discarded the targets which were in different module from the TF.

### TF-driver score

To investigate and predict the effects of a TF on a given pathway, GO term or heart phenotype gene set, we imported an approach, termed “key driver analysis”, which estimate the functions of a key driver by enrichment analysis of the driver’s targets. The targets of 123 TFs were qualified by this approach, and we scored the results by the following formula,

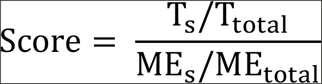

T_s_: TF targets in the pathway; T_total_: total TF’s targets; ME_s_: the over-representing module genes in the pathway; ME_total_: total module genes

For each TF, we first investigated its module and the pathways, GO terms and gene sets enriched in this module, then performed the enrichment analysis of this TF’s targets, finally selected the overlap between the two enrichment results. After determining the pathways, GO terms and gene sets, we detected TF-driver scores of individual TFs to explore their potential functions.

DNA motif enrichment analysis. To obtain regulatory regions of mouse genes, we downloaded murine heart E10.5 H3K27ac ChIP-seq data from the ENCODE project (DunhamKundaje et al., 2012) (ENCSR582SPN). The peaks, corresponding to either promoters or enhances, from −50 kb to +50 kb of the transcription start site of a gene was used for motif analysis. The enrichment of TF binding motifs were performed using the AME 4.12.0 (McLeay & Bailey, 2010) of MEME suite (FDR < 0.05). Motif visualization and similarity analysis were carried out using package motifStack 1.22.0 (Ou, Wolfe et al., 2018) and MotIV 1.34.0 (Mercier & Gottardo, 2014) in R.

### Animals

Wild type CD1 mice were obtained from Charles River Laboratories. All mouse experiments were performed according to the guideline of the National Institute of Health and the protocol approved by the Institutional Animal Care and Use Committee of Albert Einstein College of Medicine. Noontime on the day of detecting vaginal plugs was designated as E0.5.

### RNA extraction and qPCR

The hearts were isolated from E9.5 and E10.5 embryos and separated into three parts including atrioventricular canal (AVC), outflow tract (OFT) and ventricle. Tissues from four hearts were pooled as one sample and subjected to total RNAs extraction using Trizol (Life technology) according to the manufacture’s manual. First strand cDNA was synthesized using the Superscript IV Reverse Transcriptase Kit (Life technology). qPCR was performed using the Power SYBR Green PCR Master Mix (Life technology). Gene specific primers were used (Table S8). The relative expression level of genes was normalized to the expression level of Gapdh and calculated using the 2-ΔΔCTmethod. Biological repeats were performed using four different samples. Student’s t-Test was used for comparison between groups and the probability (p) value <0.05 was considered as significant.

## Acknowledgements

We would like to thank the groups of Sean M. Wu and Christine E. Seidman for sharing the scRNA-seq data, and Sean M. WU provided the information of their single-cell types.

## Author contributions

YL and DZ conceived and designed the research. YL performed the computational analysis. PL, YW, BM and BZ conducted the PCR study. YL and DZ drafted the manuscript. All authors contributed to and edited the final manuscript.

## Supplemental

Figures S1-S4 and Tables S1-S8.

## Reference

Afouda BA, Martin J, Liu F, Ciau-Uitz A, Patient R, Hoppler S (2008) GATA transcription factors integrate Wnt signalling during heart development. Development 135: 3185–90

Ahuja S, Dogra D, Stainier DYR, Reischauer S (2016) Id4 functions downstream of Bmp signaling to restrict TCF function in endocardial cells during atrioventricular valve development. Dev Biol 412: 71–82

Albert M, Schmitz SU, Kooistra SM, Malatesta M, Morales Torres C, Rekling JC, Johansen JV, Abarrategui I, Helin K (2013) The histone demethylase Jarid1b ensures faithful mouse development by protecting developmental genes from aberrant H3K4me3. PLoS Genet 9: e1003461

Anders S, Pyl PT, Huber W (2015) HTSeq—a Python framework to work with high-throughput sequencing data. Bioinformatics 31: 166–9

Bastian M, Heymann S, Jacomy M (2009) Gephi: An Open Source Software for Exploring and Manipulating Networks.

Bonilla-Claudio M, Wang J, Bai Y, Klysik E, Selever J, Martin JF (2012) Bmp signaling regulates a dose-dependent transcriptional program to control facial skeletal development. Development 139: 709–19

Carcamo-Orive I, Hoffman GE, Cundiff P, Beckmann ND, D’Souza SL, Knowles JW, Patel A, Papatsenko D, Abbasi F, Reaven GM, Whalen S, Lee P, Shahbazi M, Henrion MYR, Zhu K, Wang S, Roussos P, Schadt EE, Pandey G, Chang R et al. (2017) Analysis of Transcriptional Variability in a Large Human iPSC Library Reveals Genetic and Non-genetic Determinants of Heterogeneity. Cell Stem Cell 20: 518–532 e9

Carvalho BS, Irizarry RA (2010) A framework for oligonucleotide microarray preprocessing. Bioinformatics 26: 2363–7

Chamberlain AA, Lin MY, Lister RL, Maslov AA, Wang YD, Suzuki M, Wu BR, Greally JM, Zheng DY, Zhou B (2014) DNA Methylation is Developmentally Regulated for Genes Essential for Cardiogenesis. Journal of the American Heart Association 3

Chen J, Bardes EE, Aronow BJ, Jegga AG (2009) ToppGene Suite for gene list enrichment analysis and candidate gene prioritization. Nucleic Acids Res 37: W305–11

Chen X, Chakravarty T, Zhang Y, Li X, Zhong JF, Wang C (2016) Single-cell transcriptome and epigenomic reprogramming of cardiomyocyte-derived cardiac progenitor cells. Sci Data 3: 160079

Chen X, Qin L, Liu Z, Liao L, Martin JF, Xu J (2015) Knockout of SRC-1 and SRC-3 in Mice Decreases Cardiomyocyte Proliferation and Causes a Noncompaction Cardiomyopathy Phenotype. Int J Biol Sci 11: 1056–72

Chung S, Dzeja PP, Faustino RS, Perez-Terzic C, Behfar A, Terzic A (2007) Mitochondrial oxidative metabolism is required for the cardiac differentiation of stem cells. Nat Clin Pract Cardiovasc Med 4 Suppl 1: S60–7

Collins FS, Finnell RH, Rossant J, Wurst W (2007) A new partner for the international knockout mouse consortium. Cell 129: 235

Cunnington RH, Nazari M, Dixon IM (2009) c-Ski, Smurf2, and Arkadia as regulators of TGF-beta signaling: new targets for managing myofibroblast function and cardiac fibrosis. Can J Physiol Pharmacol 87: 764–72

da Rosa MS, Seminotti B, Ribeiro CA, Parmeggiani B, Grings M, Wajner M, Leipnitz G (2016) 3-Hydroxy-3-methylglutaric and 3-methylglutaric acids impair redox status and energy production and transfer in rat heart: relevance for the pathophysiology of cardiac dysfunction in 3-hydroxy-3-methylglutaryl-coenzyme A lyase deficiency. Free Radic Res 50: 997–1010

de la Pompa JL, Epstein JA (2012) Coordinating tissue interactions: Notch signaling in cardiac development and disease. Dev Cell 22: 244–54

DeLaughter DM, Bick AG, Wakimoto H, McKean D, Gorham JM, Kathiriya IS, Hinson JT, Homsy J, Gray J, Pu W, Bruneau BG, Seidman JG, Seidman CE (2016) SingleCell Resolution of Temporal Gene Expression during Heart Development. Dev Cell 39: 480–490

Dickinson ME, Flenniken AM, Ji X, Teboul L, Wong MD, White JK, Meehan TF, Weninger WJ, Westerberg H, Adissu H, Baker CN, Bower L, Brown JM, Caddle LB, Chiani F, Clary D, Cleak J, Daly MJ, Denegre JM, Doe B et al. (2016) High-throughput discovery of novel developmental phenotypes. Nature 537: 508–514

Dunham I, Kundaje A, Aldred SF, Collins PJ, Davis C, Doyle F, Epstein CB, Frietze S, Harrow J, Kaul R, Khatun J, Lajoie BR, Landt SG, Lee BK, Pauli F, Rosenbloom KR, Sabo P, Safi A, Sanyal A, Shoresh N et al. (2012) An integrated encyclopedia of DNA elements in the human genome. Nature 489: 57–74

Fabregat A, Jupe S, Matthews L, Sidiropoulos K, Gillespie M, Garapati P, Haw R, Jassal B, Korninger F, May B, Milacic M, Roca CD, Rothfels K, Sevilla C, Shamovsky V, Shorser S, Varusai T, Viteri G, Weiser J, Wu G et al. (2018) The Reactome Pathway Knowledgebase. Nucleic Acids Res 46: D649–D655

Fletcher A, Raza F, Buursma J, Livingston S, Kearn D, Herndon B, Vanden Heauvel G, Baybutt R, Molteni A (2015) Pathological Changes in the Heart of the Strain CUX1 Transgenic Mice. Faseb Journal 29: LB432

Gladka MM, Molenaar B, de Ruiter H, van der Elst S, Tsui H, Versteeg D, Lacraz GPA, Huibers MMH, van Oudenaarden A, van Rooij E (2018) Single-Cell Sequencing of the Healthy and Diseased Heart Reveals Ckap4 as a New Modulator of Fibroblasts Activation. Circulation

He A, Gu F, Hu Y, Ma Q, Ye LY, Akiyama JA, Visel A, Pennacchio LA, Pu WT (2014) Dynamic GATA4 enhancers shape the chromatin landscape central to heart development and disease. Nat Commun 5: 4907

Hirose-Yotsuya L, Okamoto F, Yamakawa T, Whitson RH, Fujita-Yamaguchi Y, Itakura K (2015) Knockdown of AT-rich interaction domain (ARID) 5B gene expression induced AMPKalpha2 activation in cardiac myocytes. Biosci Trends 9: 377–85

Hollnagel A, Oehlmann V, Heymer J, Ruther U, Nordheim A (1999) Id genes are direct targets of bone morphogenetic protein induction in embryonic stem cells. J Biol Chem 274: 19838–45

Jin SC, Homsy J, Zaidi S, Lu Q, Morton S, DePalma SR, Zeng X, Qi H, Chang W, Sierant MC, Hung WC, Haider S, Zhang J, Knight J, Bjornson RD, Castaldi C, Tikhonoa IR, Bilguvar K, Mane SM, Sanders SJ et al. (2017) Contribution of rare inherited and de novo variants in 2,871 congenital heart disease probands. Nat Genet 49: 1593–1601

Jongbloed MR, Vicente-Steijn R, Douglas YL, Wisse LJ, Mori K, Yokota Y, Bartelings MM, Schalij MJ, Mahtab EA, Poelmann RE, Gittenberger-De Groot AC (2011) Expression of Id2 in the second heart field and cardiac defects in Id2 knock-out mice. Dev Dyn 240: 2561–77

Kanehisa M, Furumichi M, Tanabe M, Sato Y, Morishima K (2017) KEGG: new perspectives on genomes, pathways, diseases and drugs. Nucleic Acids Res 45: D353–D361

Kim D, Langmead B, Salzberg SL (2015) HISAT: a fast spliced aligner with low memory requirements. Nat Methods 12: 357–60

Kiselev VY, Kirschner K, Schaub MT, Andrews T, Yiu A, Chandra T, Natarajan KN, Reik W, Barahona M, Green AR, Hemberg M (2017) SC3: consensus clustering of single-cell RNA-seq data. Nat Methods 14: 483–486

Kolde R (2013) pheatmap: Pretty Heatmaps. In

Lage K, Greenway SC, Rosenfeld JA, Wakimoto H, Gorham JM, Segre AV, Roberts AE, Smoot LB, Pu WT, Pereira AC, Mesquita SM, Tommerup N, Brunak S, Ballif BC, Shaffer LG, Donahoe PK, Daly MJ, Seidman JG, Seidman CE, Larsen LA (2012) Genetic and environmental risk factors in congenital heart disease functionally converge in protein networks driving heart development. Proc Natl Acad Sci U S A 109: 14035–40

Lage K, Mollgard K, Greenway S, Wakimoto H, Gorham JM, Workman CT, Bendsen E, Hansen NT, Rigina O, Roque FS, Wiese C, Christoffels VM, Roberts AE, Smoot LB, Pu WT, Donahoe PK, Tommerup N, Brunak S, Seidman CE, Seidman JG et al. (2010) Dissecting spatio-temporal protein networks driving human heart development and related disorders. Mol Syst Biol 6: 381

Langfelder P, Horvath S (2008) WGCNA: an R package for weighted correlation network analysis. BMC Bioinformatics 9: 559

Langfelder P, Luo R, Oldham MC, Horvath S (2011) Is my network module preserved and reproducible? PLoS Comput Biol 7: e1001057

Li G, Xu A, Sim S, Priest JR, Tian X, Khan T, Quertermous T, Zhou B, Tsao PS, Quake SR, Wu SM (2016) Transcriptomic Profiling Maps Anatomically Patterned Subpopulations among Single Embryonic Cardiac Cells. Dev Cell 39: 491–507

Li H, Handsaker B, Wysoker A, Fennell T, Ruan J, Homer N, Marth G, Abecasis G, Durbin R, Genome Project Data Processing S (2009) The Sequence Alignment/Map format and SAMtools. Bioinformatics 25: 2078–9

Liu Y, Morley M, Brandimarto J, Hannenhalli S, Hu Y, Ashley EA, Tang WH, Moravec CS, Margulies KB, Cappola TP, Li M, consortium MA (2015) RNA-Seq identifies novel myocardial gene expression signatures of heart failure. Genomics 105: 83-9

Liu Z, Wang L, Welch JD, Ma H, Zhou Y, Vaseghi HR, Yu S, Wall JB, Alimohamadi S, Zheng M, Yin C, Shen W, Prins JF, Liu J, Qian L (2017) Single-cell transcriptomics reconstructs fate conversion from fibroblast to cardiomyocyte. Nature 551: 100–104

Loirand G, Pacaud P (2014) Involvement of Rho GTPases and their regulators in the pathogenesis of hypertension. Small GTPases 5: 1–10

Luna-Zurita L, Stirnimann CU, Glatt S, Kaynak BL, Thomas S, Baudin F, Samee MA, He D, Small EM, Mileikovsky M, Nagy A, Holloway AK, Pollard KS, Muller CW, Bruneau BG (2016) Complex Interdependence Regulates Heterotypic Transcription Factor Distribution and Coordinates Cardiogenesis. Cell 164: 999–1014

Makinen VP, Civelek M, Meng Q, Zhang B, Zhu J, Levian C, Huan T, Segre AV, Ghosh S, Vivar J, Nikpay M, Stewart AF, Nelson CP, Willenborg C, Erdmann J, Blakenberg S, O’Donnell CJ, Marz W, Laaksonen R, Epstein SE et al. (2014) Integrative genomics reveals novel molecular pathways and gene networks for coronary artery disease. PLoS Genet 10: e1004502

McCarthy DJ, Campbell KR, Lun AT, Wills QF (2017) Scater: pre-processing, quality control, normalization and visualization of single-cell RNA-seq data in R. Bioinformatics 33: 1179–1186

McLeay RC, Bailey TL (2010) Motif Enrichment Analysis: a unified framework and an evaluation on ChIP data. BMC Bioinformatics 11: 165

Mercier E, Gottardo R (2014) MotIV: Motif Identification and Validation.

Monnier V, Iche-Torres M, Rera M, Contremoulins V, Guichard C, Lalevee N, Tricoire H, Perrin L (2012) dJun and Vri/dNFIL3 are major regulators of cardiac aging in Drosophila. PLoS Genet 8: e1003081

Muller-Borer B, Esch G, Aldina R, Woon W, Fox R, Bursac N, Hiller S, Maeda N, Shepherd N, Jin JP, Hutson M, Anderson P, Kirby ML, Malouf NN (2012) Calcium dependent CAMTA1 in adult stem cell commitment to a myocardial lineage. PLoS One 7: e38454

Naito AT, Akazawa H, Takano H, Minamino T, Nagai T, Aburatani H, Komuro I (2005) Phosphatidylinositol 3-kinase-Akt pathway plays a critical role in early cardiomyogenesis by regulating canonical Wnt signaling. Circ Res 97: 144–51

Nassiri M, Liu J, Kulak S, Uwiera RR, Aird WC, Ballermann BJ, Jahroudi N (2010) Repressors NFI and NFY participate in organ-specific regulation of von Willebrand factor promoter activity in transgenic mice. Arterioscler Thromb Vasc Biol 30: 1423–9

Olson EN (2006) Gene regulatory networks in the evolution and development of the heart. Science 313: 1922–7

Ou J, Wolfe SA, Brodsky MH, Zhu LJ (2018) motifStack for the analysis of transcription factor binding site evolution. Nat Methods 15: 8–9

Pinero J, Bravo A, Queralt-Rosinach N, Gutierrez-Sacristan A, Deu-Pons J, Centeno E, Garcia-Garcia J, Sanz F, Furlong LI (2017) DisGeNET: a comprehensive platform integrating information on human disease-associated genes and variants. Nucleic Acids Res 45: D833–D839

Qi H, Yu L, Zhou X, Kitaygorodsky A, Wynn J, Zhu N, Aspelund G, Lim FY, Crombleholme T, Cusick R, Azarow K, Danko ME, Chung D, Warner BW, Mychaliska GB, Potoka D, Wagner AJ, ElFiky M, Nickerson DA, Bamshad MJ et al. (2017) Genetic analysis of de novo variants reveals sex differences in complex and isolated congenital diaphragmatic hernia and indicates <*MYRF*> as a candidate gene. bioRxiv

Quaranta R, Fell J, Ruhle F, Rao J, Piccini I, Arauzo-Bravo MJ, Verkerk AO, Stoll M, Greber B (2018) Revised roles of ISL1 in a hES cell-based model of human heart chamber specification. Elife 7: e31706

Ritchie ME, Phipson B, Wu D, Hu Y, Law CW, Shi W, Smyth GK (2015) limma powers differential expression analyses for RNA-sequencing and microarray studies. Nucleic Acids Res 43: e47

Robinson MD, McCarthy DJ, Smyth GK (2010) edgeR: a Bioconductor package for differential expression analysis of digital gene expression data. Bioinformatics 26: 139–40

Rochais F, Dandonneau M, Mesbah K, Jarry T, Mattei MG, Kelly RG (2009) Hes1 is expressed in the second heart field and is required for outflow tract development. PLoS One 4: e6267

Russell R, Ilg M, Lin Q, Wu G, Lechel A, Bergmann W, Eiseler T, Linta L, Kumar PP, Klingenstein M, Adachi K, Hohwieler M, Sakk O, Raab S, Moon A, Zenke M, Seufferlein T, Scholer HR, Illing A, Liebau S et al. (2015) A Dynamic Role of TBX3 in the Pluripotency Circuitry. Stem Cell Reports 5: 1155–1170

See K, Tan WLW, Lim EH, Tiang Z, Lee LT, Li PYQ, Luu TDA, Ackers-Johnson M, Foo RS (2017) Single cardiomyocyte nuclear transcriptomes reveal a lincRNA-regulated de-differentiation and cell cycle stress-response in vivo. Nat Commun 8: 225

Seneviratne AN, Edsfeldt A, Cole JE, Kassiteridi C, Swart M, Park I, Green P, Khoyratty T, Saliba D, Goddard ME, Sansom SN, Goncalves I, Krams R, Udalova IA, Monaco C (2017) Interferon Regulatory Factor 5 Controls Necrotic Core Formation in Atherosclerotic Lesions by Impairing Efferocytosis. Circulation 136: 1140–1154

Sereti K-I, Nguyen NB, Kamran P, Zhao P, Ranjbarvaziri S, Park S, Sabri S, Engel JL, Sung K, Kulkarni RP, Ding Y, Hsiai TK, Plath K, Ernst J, Sahoo D, Mikkola HKA, Iruela-Arispe ML, Ardehali R (2018) Analysis of cardiomyocyte clonal expansion during mouse heart development and injury. Nature Communications 9: 754

Shannon P, Markiel A, Ozier O, Baliga NS, Wang JT, Ramage D, Amin N, Schwikowski B, Ideker T (2003) Cytoscape: a software environment for integrated models of biomolecular interaction networks. Genome Res 13: 2498–504

Shekhar A, Lin X, Liu FY, Zhang J, Mo H, Bastarache L, Denny JC, Cox NJ, Delmar M, Roden DM, Fishman GI, Park DS (2016) Transcription factor ETV1 is essential for rapid conduction in the heart. J Clin Invest 126: 4444–4459

Shirvani SM, Mookanamparambil L, Ramoni MF, Chin MT (2007) Transcription factor CHF1/Hey2 regulates the global transcriptional response to platelet-derived growth factor in vascular smooth muscle cells. Physiol Genomics 30: 61–8

Shu L, Zhao Y, Kurt Z, Byars SG, Tukiainen T, Kettunen J, Orozco LD, Pellegrini M, Lusis AJ, Ripatti S, Zhang B, Inouye M, Makinen VP, Yang X (2016) Mergeomics: multidimensional data integration to identify pathogenic perturbations to biological systems. BMC Genomics 17: 874

Sifrim A, Hitz M-P, Wilsdon A, Breckpot J, Turki SHA, Thienpont B, McRae J, Fitzgerald TW, Singh T, Swaminathan GJ, Prigmore E, Rajan D, Abdul-Khaliq H, Banka S, Bauer UMM, Bentham J, Berger F, Bhattacharya S, Bu’Lock F, Canham N et al. (2016) Distinct genetic architectures for syndromic and nonsyndromic congenital heart defects identified by exome sequencing. Nature Genetics 48: 1060

Skelly DA, Squiers GT, McLellan MA, Bolisetty MT, Robson P, Rosenthal NA, Pinto AR (2018) Single-Cell Transcriptional Profiling Reveals Cellular Diversity and Intercommunication in the Mouse Heart. Cell Rep 22: 600–610

Song X, Gao X, Lu J, Liang H, Su P, Li Q, Pang Y (2018) High mobility group box transcription factor 1 (HBP1) from Lampetra japonica affects cell cycle regulation. Dev Growth Differ 60: 146–157

Subramanian A, Tamayo P, Mootha VK, Mukherjee S, Ebert BL, Gillette MA, Paulovich A, Pomeroy SL, Golub TR, Lander ES, Mesirov JP (2005) Gene set enrichment analysis: a knowledge-based approach for interpreting genome-wide expression profiles. Proc Natl Acad Sci U S A 102: 15545–50

Takeuchi JK, Ohgi M, Koshiba-Takeuchi K, Shiratori H, Sakaki I, Ogura K, Saijoh Y, Ogura T (2003) <*Tbx5*> specifies the left/right ventricles and ventricular septum position during cardiogenesis. Development 130: 5953

Taniguchi T, Maruyama N, Ogata T, Kasahara T, Nakanishi N, Miyagawa K, Naito D, Hamaoka T, Nishi M, Matoba S, Ueyama T (2016) PTRF/Cavin-1 Deficiency Causes Cardiac Dysfunction Accompanied by Cardiomyocyte Hypertrophy and Cardiac Fibrosis. PLoS One 11: e0162513

The Gene Ontology C (2017) Expansion of the Gene Ontology knowledgebase and resources. Nucleic Acids Res 45: D331–D338

Theis JL, Sharpe KM, Matsumoto ME, Chai HS, Nair AA, Theis JD, de Andrade M, Wieben ED, Michels VV, Olson TM (2011) Homozygosity mapping and exome sequencing reveal GATAD1 mutation in autosomal recessive dilated cardiomyopathy. Circ Cardiovasc Genet 4: 585–94

Trapnell C, Cacchiarelli D, Grimsby J, Pokharel P, Li S, Morse M, Lennon NJ, Livak KJ, Mikkelsen TS, Rinn JL (2014) The dynamics and regulators of cell fate decisions are revealed by pseudotemporal ordering of single cells. Nat Biotechnol 32: 381–386

van der Maaten L, Hinton GE (2008) Visualizing High-Dimensional Data Using t-SNE. Journal of Machine Learning Research 15: 3221–3245

Waardenberg AJ, Ramialison M, Bouveret R, Harvey RP (2014) Genetic networks governing heart development. Cold Spring Harb Perspect Med 4: a013839

Waldron L, Steimle JD, Greco TM, Gomez NC, Dorr KM, Kweon J, Temple B, Yang XH, Wilczewski CM, Davis IJ, Cristea IM, Moskowitz IP, Conlon FL (2016) The Cardiac TBX5 Interactome Reveals a Chromatin Remodeling Network Essential for Cardiac Septation. Dev Cell 36: 262–75

Walkowska A, Pawlak M, Jane S, Kompanowska-Jezierska E, Wilanowski T (2016) Effects of high and low sodium diet on blood pressure and heart rate in mice lacking the functional Grainyhead-like 1 gene.

Wang RN, Green J, Wang Z, Deng Y, Qiao M, Peabody M, Zhang Q, Ye J, Yan Z, Denduluri S, Idowu O, Li M, Shen C, Hu A, Haydon RC, Kang R, Mok J, Lee MJ, Luu HL, Shi LL (2014) Bone Morphogenetic Protein (BMP) signaling in development and human diseases. Genes Dis 1: 87–105

Wickham H (2016) ggplot2: Elegant Graphics for Data Analysis. Springer-Verlag New York,

Wickham H, Francois R, Henry L, Muller K (2017) dplyr: A Grammar of Data Manipulation. In

Wickham H, Henry L (2018) tidyr: Easily Tidy Data with ‘spread()’ and ‘gather()’ Functions. In

Wu SP, Kao CY, Wang L, Creighton CJ, Yang J, Donti TR, Harmancey R, Vasquez HG, Graham BH, Bellen HJ, Taegtmeyer H, Chang CP, Tsai MJ, Tsai SY (2015) Increased COUP-TFII expression in adult hearts induces mitochondrial dysfunction resulting in heart failure. Nat Commun 6: 8245

Yaniz-Galende E, Roux M, Nadaud S, Mougenot N, Bouvet M, Claude O, Lebreton G, Blanc C, Pinet F, Atassi F, Perret C, Dierick F, Dussaud S, Leprince P, Trégouät D-A, Marazzi G, Sassoon D, Hulot J-S (2017) Fibrogenic Potential of PW1/Peg3 Expressing Cardiac Stem Cells. Journal of the American College of Cardiology 70: 728–741

Zhang B, Gaiteri C, Bodea LG, Wang Z, McElwee J, Podtelezhnikov AA, Zhang C, Xie T, Tran L, Dobrin R, Fluder E, Clurman B, Melquist S, Narayanan M, Suver C, Shah H, Mahajan M, Gillis T, Mysore J, MacDonald ME et al. (2013) Integrated systems approach identifies genetic nodes and networks in late-onset Alzheimer’s disease. Cell 153: 707–20

Zhang D, Wu B, Wang P, Wang Y, Lu P, Nechiporuk T, Floss T, Greally JM, Zheng D, Zhou B (2017) Non-CpG methylation by DNMT3B facilitates REST binding and gene silencing in developing mouse hearts. Nucleic Acids Res 45: 3102–3115

Zhang L, Nomura-Kitabayashi A, Sultana N, Cai W, Cai X, Moon AM, Cai CL (2014) Mesodermal Nkx2.5 is necessary and sufficient for early second heart field development. Dev Biol 390: 68–79

Zhang QJ, Liu ZP (2015) Histone methylations in heart development, congenital and adult heart diseases. Epigenomics 7: 321–30

Zheng GX, Terry JM, Belgrader P, Ryvkin P, Bent ZW, Wilson R, Ziraldo SB, Wheeler TD, McDermott GP, Zhu J, Gregory MT, Shuga J, Montesclaros L, Underwood JG, Masquelier DA, Nishimura SY, Schnall-Levin M, Wyatt PW, Hindson CM, Bharadwaj R et al. (2017) Massively parallel digital transcriptional profiling of single cells. Nat Commun 8: 14049

